# Unique Neurophysiological Vulnerability of Visual Attention Networks to Acute Inflammation

**DOI:** 10.1101/599159

**Authors:** Leonie JT Balter, Jos A Bosch, Sarah Aldred, Mark T Drayson, Jet JCS Veldhuijzen van Zanten, Suzanne Higgs, Jane E Raymond, Ali Mazaheri

**Affiliations:** School of Psychology, University of Birmingham, Birmingham, B15 2TT, UK; School of Sport, Exercise, and Rehabilitation Sciences, University of Birmingham, Birmingham, B15 2TT, UK; Institute of Immunity and Immunotherapy, University of Birmingham, Birmingham, B15 2TT, UK; Psychology Department, Clinical Psychology, University of Amsterdam, Amsterdam, 1018 WT, NL

**Keywords:** mild inflammation, typhoid vaccination, attention, neurophysiology, EEG

## Abstract

Illness is often accompanied by perceived cognitive sluggishness, a symptom that may stem from immune system activation. The current study used electroencephalography (EEG) to assess how inflammation affected three different distinct attentional processes: alerting, orienting and executive control. In a double-blinded placebo-controlled within-subjects design (20 healthy males, mean age = 24.5, *SD* = 3.4), Salmonella typhoid vaccination (0.025 mg; Typhim Vi, SanofiPasteur) was used to induce transient mild inflammation, while a saline injection served as a placebo-control. Participants completed the Attention Network Test with concurrent EEG recorded six hours post-injection. Analyses focused on behavioral task performance and on modulation of oscillatory EEG activity in the alpha band (9-12 Hz) for alerting as well as orienting attention and frontal theta band (4-8 Hz) for executive control. Vaccination induced mild systemic inflammation, as assessed by interleukin-6 (IL-6) levels. While no behavioral task performance differences between the inflammation and placebo condition were evident, inflammation caused significant alterations to task-related brain activity. Specifically, inflammation produced greater cue-induced suppression of alpha power in the alerting aspect of attention while individual variation in the inflammatory response was significantly correlated with the degree of alpha power suppression. Notably, inflammation did not affect orienting (i.e., alpha lateralization) or executive control (i.e., frontal theta activity). These results reveal a unique neurophysiological vulnerability to acute mild inflammation of the neural network that underpins attentional alerting functions. Observed in the absence of performance decrements, these novel findings suggest that acute inflammation requires individuals to exert greater cognitive effort when preparing for a task in order to maintain adequate behavioral performance.

**Highlights:**

- Typhoid vaccination induced a transient mild inflammatory state
- Mild inflammation alters neurophysiological process associated with attention
- Mild inflammation selectively increased alerting-related alpha suppression while behavior was unaffected
- A greater inflammatory response was correlated with more alpha suppression

## INTRODUCTION

Evidence is mounting to support the contention that immune system activation (i.e., inflammation), both chronic and acute, may degrade basic cognitive function (Allison & Ditor, 2014). Common complaints of mild cognitive deficits in conditions associated with chronic inflammation (e.g., aging, obesity, kidney disease, rheumatoid arthritis, virus infection, and neurodegenerative diseases) or acute inflammation (e.g., injury or commonplace infections) include impaired concentration (Vollmer-Conna et al., 2004), cognitive sluggishness (Smith, 2012), as well as depression (Luppino et al., 2010) and anhedonia (Freed et al., 2018). However, it remains unclear how inflammation impacts specific basic brain processes such as attention. The aim of the current study was to investigate the impact of mild acute inflammation on the fundamental cognitive function of visual attention.

Visual attention refers to the capacity to prioritize relevant information from the ever changing visual sensory environment and is a critical brain function that underpins many everyday activities (Broadbent, 1966; Posner & Rothbart, 2007). Three main roles of the attention system have been identified: Preparing the brain for upcoming salient events (alerting); preparing where to look for task relevant information (orienting); and prioritizing task-relevant information (e.g., roads signs) over concurrent, compelling but irrelevant distractions (executive control). These distinct, yet interacting, functions have been previously assessed using the Attention Network Test (ANT) (Fan et al., 2007; Fan, McCandliss, Sommer, Raz, & Posner, 2002; Posner & Rothbart, 2007).

In the current study, we utilized a variant of the ANT (Figure 1) to investigate inflammation-induced changes to the attention system. The ANT requires participants to respond to different types of cue displays that are each quickly followed by a standard target array. The latter contains a target to which a fast, accurate manual response must be made. Different cue types provide different information that can then be used by the brain to prepare for the target array. Effective use of cues is reflected by changes in brain activity that can be measured electrophysiologically during the cue-target interval and by comparing speed and accuracy of different cue-target conditions (Fan et al., 2007; Fan et al., 2002; Posner & Rothbart, 2007).

Evidence that inflammation may cause degradation of cognitive processes stems from two broad lines of enquiry. First, correlational analyses examining cognitive performance in populations with chronic inflammation (e.g., elderly, overweight, or patients with chronic inflammatory states due to disease or disorder) have generally revealed negative correlations between inflammation and cognitive performance (Lin et al., 2018; Marsland et al., 2006; but Singh-Manoux et al., 2014). However, specific evidence for inflammation-related impairment of visual attention is scant (but see Kurella Tamura et al., 2017). Moreover, these correlational studies only address effects of chronic inflammation and cannot identify inflammation as the cause of cognitive deficits, except perhaps in the rare case where appropriate mediation analyses are used (Bourassa & Sbarra, 2016). Complicating this picture is evidence that depression and poor cardiovascular health, conditions that are more prevalent in inflammation-associated states (Dhar & Barton, 2016), may themselves contribute to reduced cognitive function (e.g., Deckers et al., 2017).

A second line of research linking inflammation and cognitive degradation involves experimental induction of transient inflammation via administration of immune-activating agents, such as bacterial endotoxin, in otherwise healthy participants. However, previous studies using endotoxin induced inflammation reported no effects on cognitive tests presumed to evaluate attention processes (Grigoleit et al., 2010; Krabbe et al., 2005; Reichenberg et al., 2001; van den Boogaard et al., 2010). Similarly, Brydon et al., (2008) reported no effect on putative attention measures using vaccination against *Salmonella typhi* as a low-grade inflammatory stimulus. Interestingly, while performance was not affected, this study showed compared to a placebo condition, greater BOLD activity during task performance, perhaps reflecting that increased effort was needed. Moreover, these aforementioned studies used coarse cognitive tests (e.g., digit span forward, digit symbol test, color-word Stroop test) that more likely index memory, learning and other high-level executive functions, leaving open the question of whether inflammation disrupts the functions that support visual attention, *per se*. Furthermore, previous work has focused almost exclusively on behavioral measures of attention, leaving largely unexplored the effects of acute inflammation on the underlying neurophysiological mechanisms. However, knowledge of these effects is of value for at least two reasons. First, absence of behavioral effects does not imply the absence of underlying neurophysiological effects of inflammation. Second, understanding neurophysiological mechanisms that underpin inflammation-associated cognitive changes may open up possibilities for early markers for those at risk to develop cognitive dysfunction. Experimental studies of inflammation have shown that induced inflammation induced by means of interferon-alpha, typhoid vaccination or endotoxin was associated with increased task-relevant neural responses, while behavioral performance was unaffected (Brydon et al., 2008; Capuron et al., 2005; Kullmann et al., 2013). This combination of increased neural recruitment and preserved behavioral performance has been interpreted as reflecting increased effort needed to perform the task.

To address effects of inflammation on attention, the current study experimentally induced acute mild inflammation and assessed visual attention using the ANT paradigm. Typhoid vaccination was used to induce mild inflammation without the concurrently inducting fever and flu-like symptoms as typically occurs with endotoxin (e.g., nausea, headache and extreme fatigue; e.g., Harrison et al., 2015) that could directly degrade cognition (Grigoleit et al., 2011; Lasselin et al., 2016). We used a randomized double-blind crossover design with a saline injection as the placebo condition. The analysis focused on injection condition-dependent changes in oscillatory activity in the electroencephalogram (EEG) alpha band to the onset of visual alerting cues, a measure reflecting mental preparation effort (Fink, Grabner, Neuper, & Neubauer, 2005; Keil, Mussweiler, & Epstude, 2006; Sawaki, Luck, & Raymond, 2015). Frontal-midline theta-band oscillations were assessed as a measure for executive control (Cavanagh & Frank, 2014). Activity in the theta band has been associated with aspects of task monitoring and error detection, that is linked to executive attention (Fan et al., 2007). According to the inhibition-timing hypothesis (Klimesch, Sauseng, & Hanslmayr, 2007), alpha selectively increases in a region that is task-irrelevant; when attention is being cued to the left or right visual hemifield, higher occipital alpha power is measured contralateral to the unattended side than contralateral to the attended side. The ratio between left and right occipital alpha, referred to as the Alpha Lateralization Index (ALI), was assessed as an index of the efficiency of orienting attention (Haegens, Handel, & Jensen, 2011). The primary prediction was that vaccine-induced inflammation would degrade the brain’s ability to prepare for upcoming task-relevant events, resulting in either reduced behavioral indices of alerting functions and/or neurophysiological evidence of enhanced preparatory effort allowing behavioral performance to be maintained.

## METHODS & MATERIALS

### Participants

Twenty healthy young male students from the University of Birmingham (*M* age = 24.5, SD = 3.4 years) were enrolled via advertisement and participated in exchange for course credits or £40. Individuals were excluded if they reported being a smoker, having a history of or suspected vaccine-related allergy, food allergy or intolerance, inflammatory, cardiovascular, neurological, mental health or immune-related disorder, visual impairment (unless corrected to normal), and those on any medication 7 days prior to the test days. Mean body mass index (BMI) was 24.5 (SD = 3.4), 16.6 – 29.2 kg/m^2^. The study was conducted according to the guidelines laid down in the Declaration of Helsinki and all procedures were approved by the local Research Ethics Committee of the National Health Service (NHS).

### General procedures

Participants visited the behavioral immunology laboratory on three separate occasions: one familiarization session, followed by two separate test days (i.e., receiving vaccination or saline placebo, randomly assigned) at least one week apart. The study was carried out in a double-blind placebo-controlled crossover fashion. On arrival in the morning, a blood sample was taken before participants received the vaccination or placebo injection. After injection, participants had a 4-hour break followed by a standardized lunch and EEG set-up preparations. A second blood sample was taken about 5h30 post-injection; then, the ANT was completed while EEG recordings took place (about 6 hours post-injection), followed by a set of other cognitive tests. Other cognitive tests, not reported here, included measures of emotion recognition, memory, learning, and response inhibition. For a detailed description of the complete study procedures, see Balter et al. (2018). The final blood sample was taken about 8 hours post-injection. Mood and sickness symptoms and tympanic body temperature were assessed at several intervals during the test day, including before injection, at 5h30m, and at 8 hours post-injection. Test timings were identical across visits, as were the procedures, except for the type of injection (vaccine or saline placebo).

### Typhoid vaccination

Typhoid vaccine was selected as a low-grade inflammatory stimulus as this vaccine is known to induce increases in circulating pro-inflammatory cytokine levels without inducing significant effects on sickness symptoms such as fever (e.g., Paine, Ring, Bosch, Drayson, & Veldhuijzen van Zanten, 2013). Participants received 0.5 mL Salmonella typhi capsular polysaccharide vaccine (0.025 mg in 0.5 mL, Typhim Vi, SanofiPasteur, UK) or a saline placebo (0.5 mL) via intra-muscular injection in the deltoid muscle of the non-dominant arm by a certified nurse on each test day.

### Mood and sickness symptom assessment

Current mood and presence of sickness symptoms was assessed using a modified version of the Profile of Mood States – Short Form (POMS-SF; Curran, Andrykowski, and Studts 1995). The version used here comprised 38 items each beginning with ‘How are you feeling right now’ followed by a word descriptive of one of eight states (tension-anxiety, anger-hostility, fatigue-inertia, vigor-activity, confusion-bewilderment, depression-dejection, physically well - physically ill, withdrawn-sociable). Ratings were made using a five-point Likert scale (0 = not at all, 1 = a little, 2 = moderately, 3 = quite a bit, to 4 = extremely). Scores for POMS subscales were computed by summing ratings on individual items.

### Blood sampling

Blood (6 ml) was collected into a vacutainer containing ethylenediaminetetraacetic acid (EDTA) as anticoagulant (Becton Dickinson Diagnostics, Oxford, United Kingdom). Samples were immediately centrifuged at 1500 *g* for 10 min at 4 °C and plasma was aliquoted and stored at −80 °C for later cytokine assessment of plasma interleukin-6 (IL-6). Plasma IL-6 was assessed in duplicate using high-sensitivity enzyme-linked immunosorbent assay (ELISA; Quantikine HS Human IL-6 ELISA, R&D Systems, UK) in accordance with the manufacturer’s instructions. The limit of detection of this assay was 0.11 pg/mL, with an intra-assay coefficient of variation of 4.2%. All samples from the same participant were assayed in the same run. Assessment of IL-6 primarily acted as an inflammation check and no causal assumptions about the role of IL-6 in the observed effects can be made.

### Behavioral and Electrophysiological data acquisition

Participants were seated at approximately 80 cm distance in front of a high-resolution 17’ LCD color monitor. E-Prime 2.0 (Psychology Software tools, Inc., Sharpsburg, PA, USA) was used to present stimuli and record behavioral data. Task responses were recorded via a standard computer keyboard. Electroencephalograms were obtained in a temperature-controlled environment using a 64-channel Ag/AgCl electrode 10-10 WaveGuard cap and eego™ sports amplifier from ANT (https://www.ant-neuro.com). The data was acquired with a sampling rate of 1024 Hz with online Cpz reference. Electrodes impedance was maintained below 20 kΩ as recommended by the manufacturers. Horizontal eye movements were monitored via two bipolar electrodes placed at the external canthi of each eye.

### Attention Network Test

In the current study we utilized a lateralized variant of the ANT commonly used in previous studies (Fan et al., 2002). Each trial began with the presentation of a black central fixation cross (1° of visual angle in diameter) that remained on the screen throughout the trial. See Figure 1. After a randomly jittered interval of 400-500 ms, one of three cue conditions occurred. (1) No Cue: A blank screen except for the fixation cross until target array presentation occurred 1200 ms later, providing no information regarding when or where the target would appear; (2) Double Cue: Two asterisks (each 0.6° in diameter) presented 2.3° to the left and right of the fixation cross along the horizontal meridian for 100 ms, providing certainty over when but not where the target would appear; and (3) Spatial Cue: A single asterisk presented either 2.3° to the left or right of the fixation cross for 100 ms, signaled the target location on 100% of the trials (spatial cue) providing both temporal and spatial certainty about the target’s appearance. The target array was presented 1200 ms after cue onset and disappeared immediately upon response or within 2000 ms. The target stimulus was a central arrow (0.6° tall X 1.7° wide) presented simultaneously with four similar (flanker) arrows pointing in the same (congruent) or opposite (incongruent) direction as the target. Flankers were positioned directly above and below the target; all arrows were aligned vertically in a column (3.3° total height; arrows separated by 0.1°) and could appear 2.3° to the left or right of fixation. The participant’s task was to report the direction of the central target arrow by pressing the “k” key for up or the “m” key for down using the dominant hand as quickly and accurately as possible. Each trial lasted 2250 ms on average.

Participants performed one practice block of 32 trials followed by six experimental blocks. Within each block, each cue condition was presented 16 times in a pseudorandom order with each combination of target array location and congruency being equally likely to occur within each cue condition.

**Figure 1.**
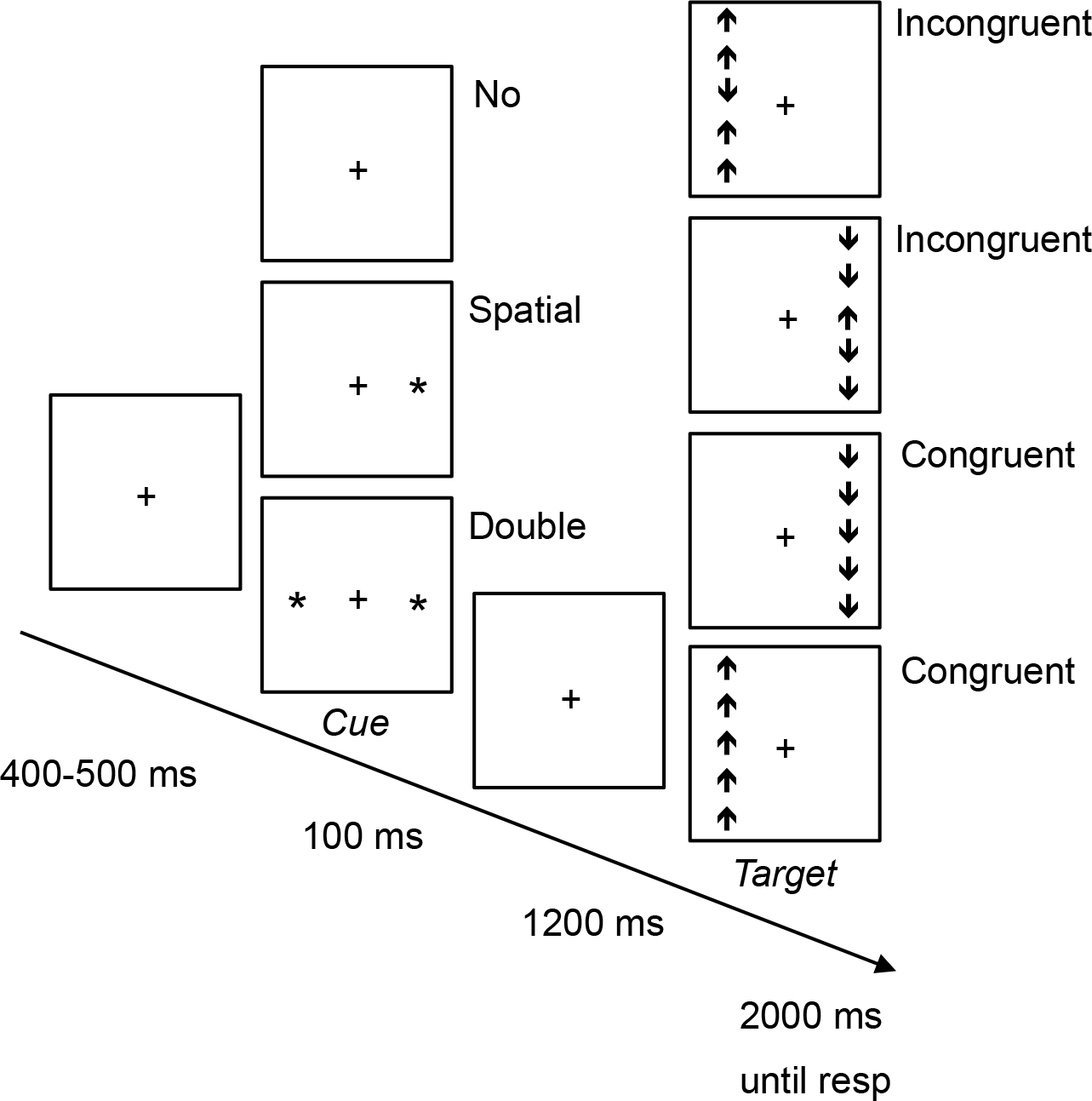
The sequence of displays presented in each trial of the lateralized ANT task. The cue display had three equally likely conditions: No Cue, Double Cue and Spatial Cue (100% valid). The target array comprised a central arrow and four identical flankers that were either congruent or incongruent with the target arrow. The task was to report the direction (up, down) of the center target arrow as quickly and accurately as possible.

In this test, differences in response time (RT) and accuracy for different cue and target flanker conditions were used to index the efficiency of two different cueing effects: (1) alerting, i.e., computing the difference between trials with a double cue and trials with no cue; and (2) orienting, i.e., computing the difference between trials with spatial cues that validly predicted the spatial location of the target and those with double cues, offering no prediction. For EEG, orienting of attention was calculated as the Alpha Lateralization Index (ALI) which is described in the next section. In addition, flanker congruency effect referred to as (3) “executive control” was determined by comparing congruent and incongruent flanker trials regardless of cue condition. Psychomotor speed was computed as the grand average RT across all cue and target-flanker conditions.

### Behavioral data analysis

Trials with incorrect responses or with RTs <150 ms or >1500 ms were excluded from any further analysis, accounting for 1.1% of the data. In addition, RTs >3 SDs from the mean for each combination of cue and target flanker condition for each participant were also excluded, accounting for an additional 1.2% of the data. All data for two participants was excluded from analysis as a result of equipment failure (no behavioral file was saved).

Mixed design repeated measures ANOVAs and paired samples *t*-tests were used where appropriate. We analyzed both RT and accuracy for testing order effects (Day 1 versus Day 2) and found neither main nor interaction effects (all F’s < 1). To assess whether the inflammatory response was associated with measures of interest (e.g., behavioral measures, alpha power, frontal theta power, ALI), bivariate correlational analyses were performed with the inflammatory response to the vaccine. The inflammatory response was defined as the IL-6 level at 5h30 post-injection in the vaccine condition minus the IL-6 level at 5h30 post-injection in the placebo condition. In addition to traditional null hypothesis significance testing, Bayes Factors were calculated for non-significant effects via Bayesian ANOVAs and correlation analyses using default prior probabilities in JASP version 0.9. Bayes factors provide relative evidence of both the null (H_0_) and alternative hypothesis (H_A_), compared to the conclusions about the null hypothesis proffered by traditional null hypothesis significance testing. To allow for clear interpretation, the approximate classification scheme of Wagenmakers et al., (2017) was used which states that an estimated Bayes Factor (BF_10_; H_0_/ H_A_) value <1 supports evidence in favor of H_0_. For example, a BF_10_ of .25 indicates that the H_0_ is 4 times (1:.25) more likely than the H_A_. Values close to 1 are not informative and a BF_10_ between 1 and 3 provides anecdotal evidence for the H_A_. A BF_10_ between 1 and .33 provides anecdotal evidence for the H_0_ (e.g., 1:3 probability in favor of H_0_) and a BF_10_
between 0.33 and .10 provides moderate evidence for H_0_. A BF_10_ < 0.10 provides strong evidence for H_0_.

### Time-Frequency analysis

Offline processing and analyses were performed using the matlab toolbox EEGLAB (Delorme & Makeig, 2004) and Fieldtrip (Oostenveld, Fries, Maris, & Schoffelen, 2011). Continuous EEG data were offline re-referenced to the average of all scalp electrodes, high-pass filtered at 0.1 Hz and low-pass filtered below 40 Hz using a two-way, fourth-order Butterworth filter. To remove ocular and muscle artifacts, independent components analysis (ICA) was performed using the EEGLAB toolbox (on average, 6 components per participant were removed, SD = 2.0). There was no significant difference between the number of components accepted in the vaccine versus placebo condition, *t*(19)= 1.24, *p*=.230. Next, continuous EEG data were epoched from −2000 ms to 2000 ms, time-locked to cue onset. Only correct trials were included in the analyses. Time-frequency representations of power were estimated per trial, using sliding Hanning tapers having an adaptive time window of five cycles for each frequency of interest (ΔT = 5/f). For the ‘alerting’ component of the ANT we focused on the difference in modulation of alpha power (averaged across 9-12 Hz) in the interval between 200 ms post-cue onset to target onset for double versus no-cues. We chose to analyze activity 200ms after the onset of cues as to avoid spectral contamination from the sensory evoked responses into the alpha band. The difference in alpha modulation (over time and electrodes) between ‘placebo vs vaccine’ conditions was assessed by means of cluster-based permutation procedure (Maris & Oostenveld, 2007) implemented in the Fieldtrip toolbox (Oostenveld et al., 2011). This test controls the Type I error rate involving multiple comparisons (e.g. multiple channels or time-frequency tiles). A probability value here is obtained through the Monte Carlo estimate of the permutation *p*-value of the cluster of channels by randomly swapping the labels (i.e., condition) in participants 5000 times and calculating the maximum cluster-level test statistic.

For the ‘executive control’ aspect of the ANT, we examined the post-target differences in theta power between incongruent target flankers and congruent target flankers in the interval 200-700 ms post-target onset (as to avoid spectral contamination from the sensory evoked responses. The difference in theta modulation related to congruency (over time and electrodes) between ‘placebo vs vaccine’ conditions was also assessed by means of a cluster-based permutation procedure.

Lastly, for the ‘orienting’ aspect of the ANT, the alpha lateralization index (ALI) was calculated for the left and right cue for early (0 to 500 ms post-cue) and late (500 to 1000 ms post-cue) time points. The region of interest (ROI) included posterior channels for the left (P3, P5, P7, PO5, PO7) and right hemisphere (P4, P6, P8, PO6, PO8). The ALI was calculated as follows:

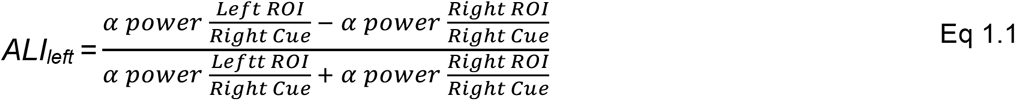

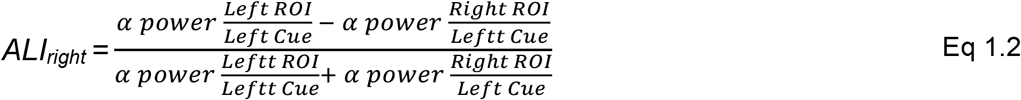

A repeated measures ANOVA with cue (left, right), time (early, late), and injection condition (placebo, vaccine) was conducted to assess the effect of injection condition on the ALI. Levene’s test of Equality of Variances and Mauchly’s Test of Sphericity were used to test for assumption violations with adjustments made using the Greenhouse-Geisser corrections; Bonferroni corrections were applied to post-hoc pairwise comparisons (two-tailed unless stated otherwise) to control for type I error rate; and alpha was set at .05.

## RESULTS

### Physiological responses

Participants showed a significantly greater peak in plasma IL-6 to typhoid vaccination (+4.0 pg/mL, *SD* =1.6) as compared to placebo (−0.1 pg/mL, *SD* = .3) (time (0h, 5h30, 8h) x injection condition), *F*(2, 26) = 30.10, *p* < .001, η_p_^2^ = .82). The peak IL-6 response (See Table1) occurred at 5h30m post-injection for most participants. IL-6 remained elevated at 8 hours post injection (+2.3 pg/mL, SD = 1.9), *t*(16) = −7.03, *p* < .001). As such from this point on we will refer to the vaccine condition as the inflammation condition for the remainder of the text. Peak body temperature was similar across conditions (inflammation *M* = 36.9°C, SD = 0.3; placebo *M* =36.8°C, SD = 0.3), confirming absence of fever.

**Table 1.** Mean (SD) interleukin-6 (IL-6) concentrations (pg/mL) separated by injection condition. Column labels represent time since injection.

|  | 0 hours | 5h30m | 8 hours |
| --- | --- | --- | --- |
| Placebo | 0.9 (0.3) | 0.8 (0.4) | 0.8 (0.4) |
| Vaccine | 1.2 (0.5) | 5.1 (1.4) | 3.9 (1.3) |

### Mood and sickness symptoms

No significant time by injection condition interactions were evident for any of the POMS subscales or total mood score (all *F*’s < 1). POMS data are summarized in Table 2.

**Table 2.** Mean (SD) POMS subscales (mood and physical and behavioral symptoms) separated by injection condition. Column labels represent time since injection.

|  | 0 hours |  | 5h30m |  | 8 hours |  |
| --- | --- | --- | --- | --- | --- | --- |
|  | Placebo | Inflammation | Placebo | Inflammation | Placebo | Inflammation |
| Anger-Hostility | 0.4 (1.0) | 0.2 (0.5) | 0.6 (1.4) | 0.2 (0.4) | 0.8 (2.0) | 1.0 (2.0) |
| Confusion-Bewilderment | 3.4 (1.6) | 3.3 (1.5) | 3.3 (1.7) | 3.1 (1.0) | 4.5 (2.1) | 4.5 (1.9) |
| Depression-Dejection | 0.4 (1.0) | 0.4 (1.0) | 0.3 (0.6) | 0.2 (0.6) | 0.8 (2.2) | 1.5 (2.6) |
| Fatigue-Inertia | 2.6 (2.9) | 2.7 (2.3) | 2.5 (3.2) | 1.8 (2.5) | 4.9 (3.2) | 5.7 (3.8) |
| Tension-Anxiety | 0.7 (1.1) | 0.6 (0.9) | 1.1 (2.2) | 1.1 (1.6) | 1.2 (2.4) | 1.0 (1.9) |
| Vigor-Activity | 6.8 (5.5) | 6.5 (3.4) | 6.6 (3.9) | 6.7 (3.5) | 3.9 (4.2) | 3.4 (3.2) |
| Withdrawn-Sociable | 2.2 (1.2) | 2.2 (1.2) | 2.1 (1.3) | 2.0 (1.0) | 2.6 (1.3) | 2.4 (1.3) |
| Physically well-III | 0.6 (1.2) | 0.7 (1.6) | 0.8 (2.0) | 0.6 (1.5) | 0.9 (1.8) | 0.7 (1.4) |
| Total mood | 0.7 (8.8) | -0.2 (5.6) | 1.3 (9.9) | -0.4 (6.2) | 8.2 (12.1) | 9.4 (11.4) |

### Behavioral attention measures

ANOVA of individual condition means revealed no significant main effect of injection condition on RT (placebo *M* = 480 ms, SE = 11; inflammation *M* = 486 ms, SE = 11) (*F*(1, 16)= 1.00, *p*=.332, η_p_^2^ = .06). However, as expected, main effects of Cue condition (*F*(2, 32) = 42, *p* <.001, η_p_^2^ = .72) and Target Flanker condition (*F*(1, 16) = 123, *p* < .001, η_p_^2^ = .89) were found. Injection condition did not interact with Cue or Target Flanker condition (F’s < 1.1). Mean RTs and accuracy for each cue and target flanker condition for the placebo and inflammation condition are shown in Table 3. Subsequently, repeated measures ANOVAs were performed for each cueing comparison. Across injection condition, RTs for No Cue trials were 20 ms (SE = 3 ms) slower as compared to Double Cue trials (alerting effect) (*F*(1,16) = 55.73, *p* < .001, η_p_^2^ = .78). RTs for Double Cue trials were 8 ms (SE = 3 ms) slower as compared to Spatial Cue trials (orienting effects) (*F*(1, 16) = 9.38, *p* < .001, η_p_^2^ = .37). Lastly, RTs for trials with incongruent target flankers were 38 ms (SE = 3 ms) slower as compared to trials with congruent target flankers (executive control effects) (*F*(1, 16) = 124.84, *p* < .001, η_p_^2^ =.89). Injection condition did not significantly affect alerting (*F*=1.71, *p*=.209), orienting (*F* =1.47, *p*=.243), or executive control (*F*=0.23, *p*=.883) performance. None of the cueing comparisons significantly correlated with the inflammatory response (*p*’s;> 0.7). Bayesian repeated measures ANOVA showed moderate (RT alerting BF_10_ = .27; RT executive control BF_10_ = .14) and anecdotal (RT orienting BF_10_ = .41) evidence for the null hypothesis (H_0_; no effect of injection condition) against the alternative hypothesis (H_A_). Furthermore, Bayesian correlation analysis showed moderate evidence in favor of H_0_ for a correlation between IL-6 and cueing effects (alerting: BF_10_ = .30; orienting: BF_10_ = .30; executive control: BF_10_ = .30). RT and accuracy cueing effects for alerting and orienting and executive control are shown in Table 4.

Similar to RT, accuracy was not significantly affected by injection condition (placebo *M* = 96.8%, SE = 0.5; inflammation *M* =97.0%, SE = 0.6) (*F*(1, 16) = 0.165, *p* = .690, η_p_^2^ =.01). A main effect of Target Flanker condition (*F*(1, 16) = 32, *p* < .001, η_p_^2^ = .67) was found and no effect of Cue condition was evident (*F*(2, 32) = 1.42, *p* = .256, η_p_^2^ = .08). Injection condition did not interact with Cue or Target Flanker condition (F’s < 1). ANOVAs were performed for each cueing comparison. No significant alerting (*F* < 1) and orienting effects (*F*(1, 16) = 3.83, *p* =.068) of accuracy were found. Across injection condition, a significant executive control effect was found (*M* = −3.8%, SE = 0.6%) (*F* (1, 16) = 35.36, *p* < .001, η_p_^2^ =.69). No effects of injection condition were evident (*F*’s < 1) and correlations between cueing comparisons and the inflammatory response were non-significant (*p*’s > 0.1). Bayesian repeated measures ANOVA and correlation analysis showed moderate (ANOVA alerting BF_10_ = .33; ANOVA executive control BF_10_ = .33) and anecdotal evidence for H_0_ against H_A_ (ANOVA orienting BF_10_ = .45; correlation alerting: BF_10_ = .77; correlation orienting: BF_10_ =.64; correlation executive control: BF_10_ = .37).

These results suggest that information of the cues was used to improve performance to the same degree in the placebo and inflammation condition.

**Table 3.** Mean RT in ms and accuracy in % (SD) as a function of target flanker condition and cue condition for placebo and inflammation condition.

|  | Flanker condition |  |  |  | Cue condition |  |  |  |  |  |  |  |
| --- | --- | --- | --- | --- | --- | --- | --- | --- | --- | --- | --- | --- |
|  | Congruent |  | Incongruent |  | No |  | Double |  | Spatial |  | Mean |  |
|  | RT | % | RT | % | RT | % | RT | % | RT | % | RT | % |
| Placebo | 459 | 98.7 | 497 | 94.7 | 493 | 96.9 | 475 | 96.0 | 465 | 97.5 | 477 | 96.7 |
|  | (41) | (1.3) | (47) | (3.8) | (47) | (2.6) | (44) | (3.3) | (44) | (2.0) | (44) | (2.2) |
| Inflammation | 468 | 98.6 | 506 | 94.7 | 502 | 96.8 | 481 | 96.1 | 474 | 97.2 | 487 | 96.6 |
|  | (44) | (1.5) | (49) | (3.2) | (48) | (3.1) | (44) | (2.8) | (44) | (2.3) | (46) | (2.1) |

**Table 4.** Mean RT in ms and accuracy in %(SD)attention effects for placebo and inflammation condition.

|  | Alerting |  | Orienting |  | Executive Control |  |
| --- | --- | --- | --- | --- | --- | --- |
|  | RT | % | RT | % | RT | % |
| Placebo | -18 (15) | 0.8 (3.0) | 10 (11) | -1.4 (2.9) | 38 (16) | -4.0 (3.5) |
| Inflammation | -21 (14) | 0.7 (3.2) | 6 (15) | -1.1 (2.3) | 37 (13) | -3.9 (2.9) |

### Inflammation extenuates alpha suppression related to ‘alerting’

The cue-induced modulation of alpha power in the alerting dimension of the ANT (i.e. Double Cue - No Cue) can be seen in Figure 2. Cue-induced alpha suppression was significantly greater in the inflammation condition relative to placebo, in the interval 200-300 ms after cue onset (*t*(16) = −46.84, *p* = .037, Monte Carlo *p*-value, corrected for multiple comparisons). This effect was most pronounced over a central, frontal and frontal-central cluster of electrodes. Individual variation in the log IL-6 response to the vaccine was significantly negatively correlated with alpha power (*r*(15) = −.536, *p* = .048) (Figure 3). Bayesian correlation analysis showed anecdotal evidence in favor of H_A_ for a negative correlation between IL-6 and alpha power (BF_10_ = 1.95).

**Figure 2.**
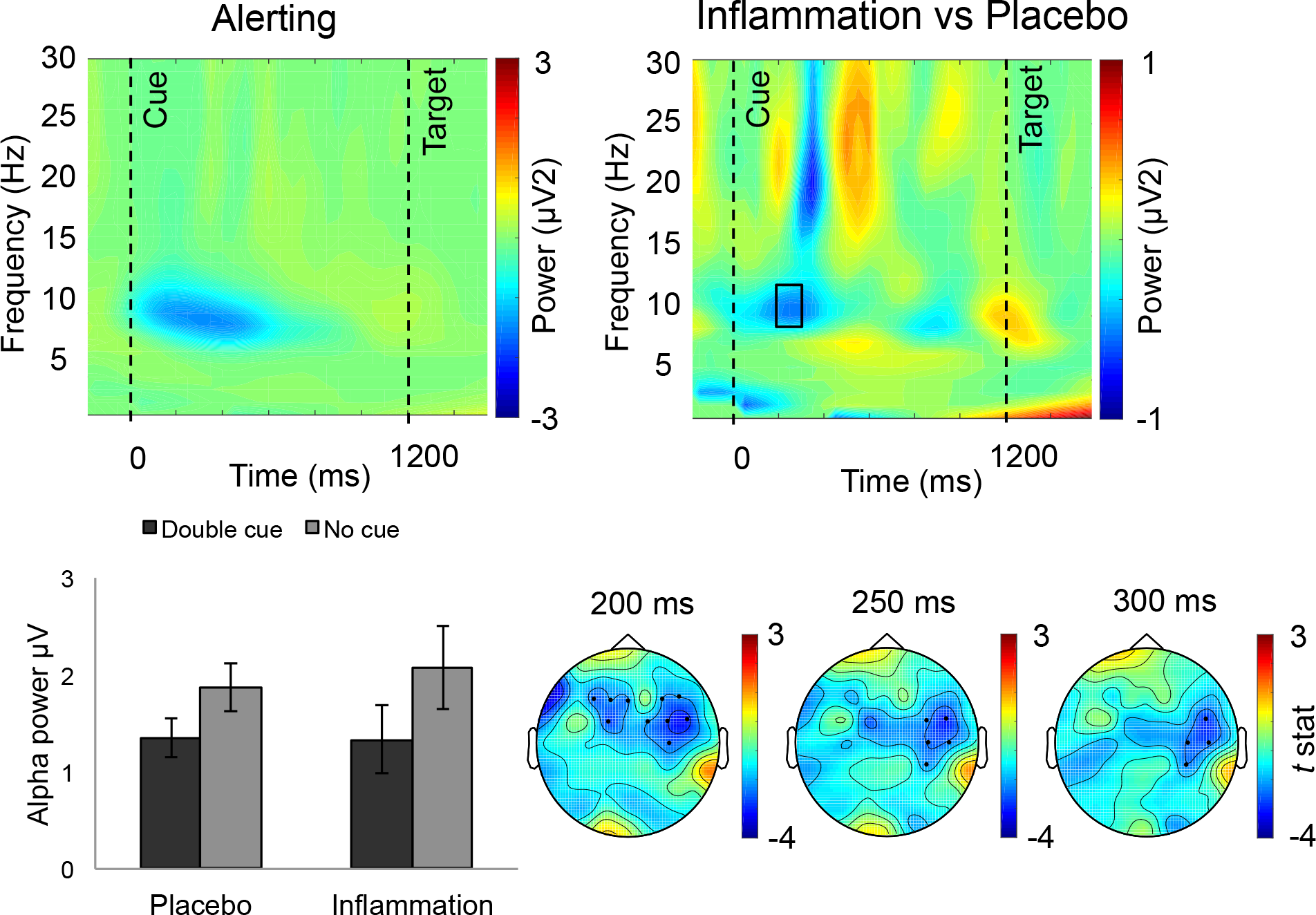
Alpha modulation related to ‘alerting’ dimension of the ANT (A) Power-locked alpha activity cued to onset of Double versus No Cue across injection conditions averaged over the significant cluster of electrodes and time interval (200-300 ms) that was revealed with random-cluster permutation tests. (B) The random-cluster permutation test revealed significantly greater alpha suppression in the inflammation compared to the placebo condition 200-300 ms after cue onset (highlighted in the black box). (C) Bar graph showing the alpha power for Double and No cue for the placebo and inflammation condition. Error bars represent standard error of the mean. (D) The dots illustrate the clusters of electrodes that show the most pronounced difference between placebo and inflammation condition over the time interval in which a significant difference was found using the random-cluster permutation test (200-300 after cue onset).

**Figure 3.**
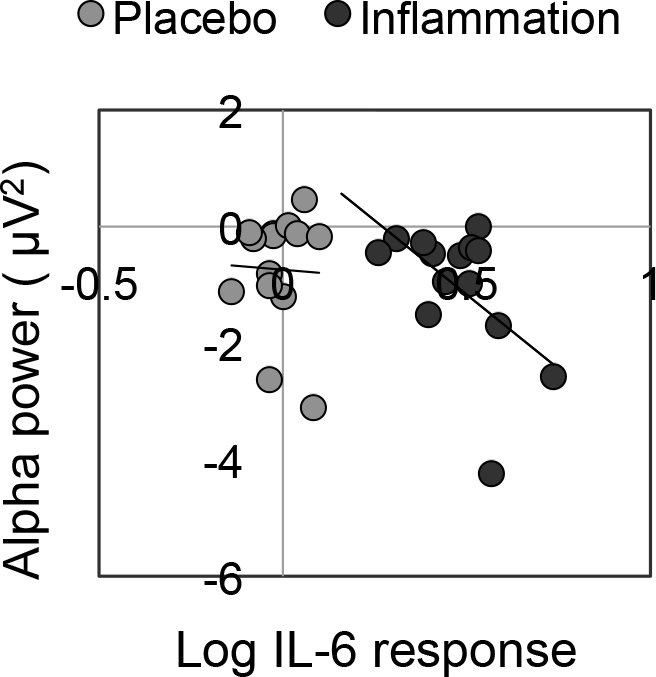
Significant negative correlation between the IL-6 response to the vaccine and alpha power to alerting cues in the inflammation conditions (black dots). Participants with a larger IL-6 response showed greater alpha power suppression. No significant correlation was found between the IL-6 response to the placebo and alpha power to altering cues in the placebo condition (gray dots). Alpha power was averaged across the electrodes (see Figure 2d) and the time period in which placebo and inflammation showed the greatest difference (200-300 ms post-cue).

### Inflammation did not significantly affect the theta increase related to executive control

While there is greater target-locked frontal theta oscillatory activity in the incongruent versus congruent target flanker condition (*t*(16) = 16.7, *p* < .001), no significant difference in theta increase between placebo and inflammation was found using cluster-based permutation analysis procedure (*t*(16) = −18.03, *p* = .433). See Figure 4. Individual variation in the IL-6 response to the vaccine was not significantly correlated with theta activity (averaged across the electrodes Cz, Cpz) in the executive control domain of attention (*r*(16) = .011, *p* =.969), with anecdotal (BF_10_ = .33) evidence in favor of H0.

**Figure 4.**
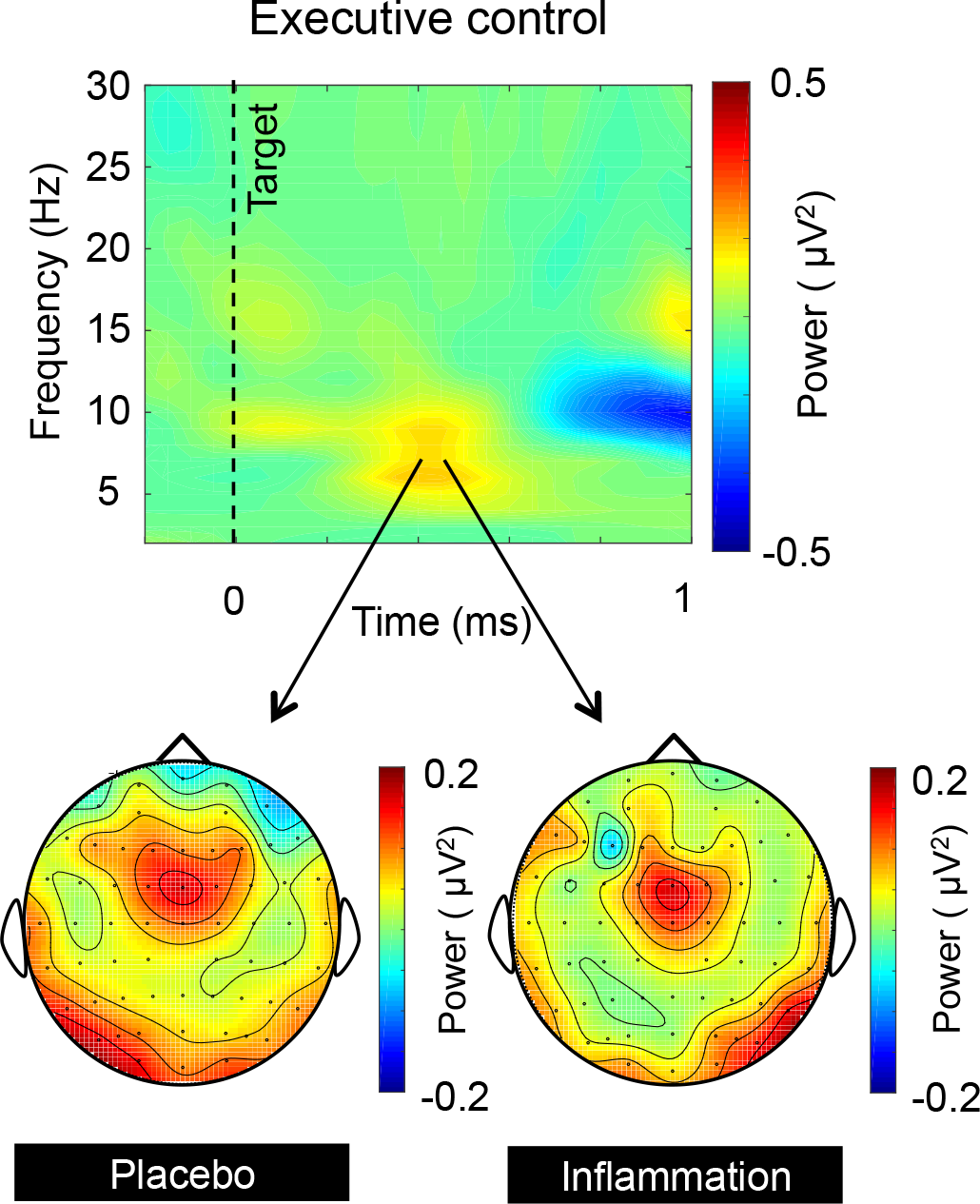
Frontal theta (Cz, Cpz) activity for executive control (A) Time-frequency representations of power-locked activity cued to the onset of targets (Incongruent-Congruent target flankers) of the ANT for placebo and inflammation condition. (B) Scalp topography of target-related theta activity. The distribution of theta power (4-7Hz) from 300 to 500 ms after target onset shown for the placebo and inflammation condition. Theta power is maximal over frontal midline electrodes.

### Inflammation did not affect the alpha lateralization related to orienting

The cue-induced alpha lateralization index (ALI) left and right for early (0-500 ms post-cue) and late (500-1000ms post-cue) processing was calculated (see Eq 1.1 and Eq 1.2) and can be seen in Figure 5. No significant effect of injection condition on the ALI (*F* = .60, *p* = .451) was found nor interactions between injection condition and cue (left, right) or time (early, late) (*F*’s < 1). A time x cue interaction revealed that, for the late time point, ALI_left_ was more negative as compared to ALI_right_ (*F* (1, 14) = 7.82, *p* = .014, η_p_^2^ = .36). The IL-6 response was not significantly correlated with the early or late ALI_left_ and ALI_right_ (*p*’s > .1). Bayesian correlation analysis showed anecdotal evidence in favor of H_0_ for a correlation between IL-6 and alpha power (Early ALI_left_ BF_10_ = 0.37; Late ALI_left_ BF_10_ = 0.35; Early ALI_right_ BF_10_ = 0.53; Early ALI_right_ BF_10_ = 0.94).

**Figure 5.**
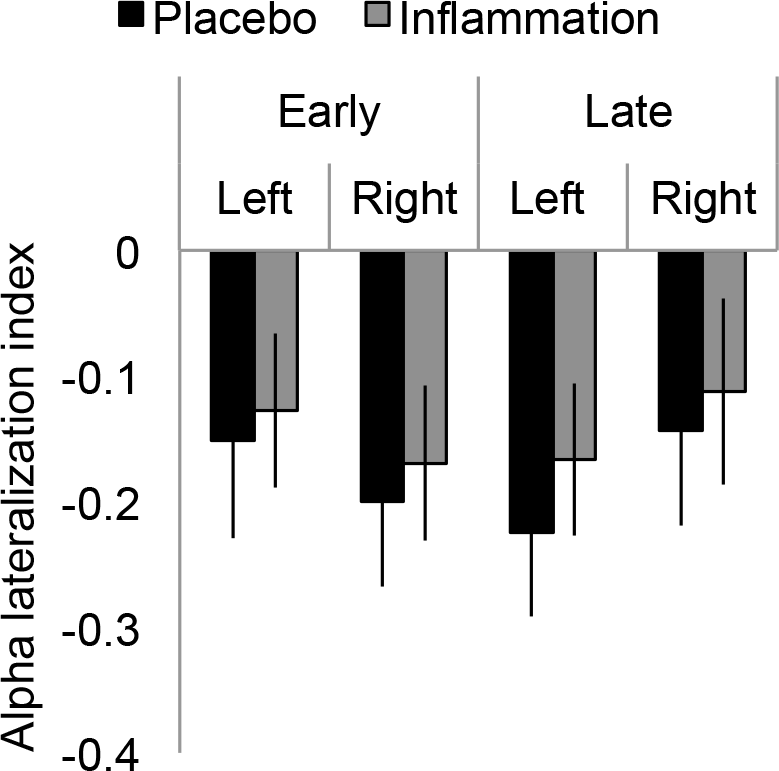
Mean Alpha Lateralization Index (ALI) 0-500 ms post-cue (early) and 500-1000 ms post-cue (late) onset for the left and right cue separately for the placebo and inflammation condition. A more positive value of ALI is indicative of a shift in alpha power to the left hemisphere, a more negative value of ALI means a shift to the right hemisphere.

## DISCUSSION

The current study used the ANT to investigate how acute low-grade inflammation affected the alerting, orienting and executive components of visual attention at a neurophysiological and behavioral level. Typhoid vaccination given to the participants effectively induced mild inflammation, as measured by IL-6, without evoking concurrent fever or flu-like symptoms, a finding in line with results from a larger cohort (Balter et al., 2018). The primary finding of the current study is that transient mild inflammation affected the alerting network, but left the other attention components, orienting and executive relatively unaffected. Specifically, we observed that the alpha suppression response to cues that provide temporal information about imminent targets stimuli was significantly more pronounced in the vaccination condition than the placebo. Importantly, this alteration in the brain’s preparation for a target display did not result in any overt behavioral change in performance. The results demonstrate for the first time that a sub-process of attention, i.e., alerting, is selectively vulnerable to the effects of acute mild inflammation.

The degree to which alpha power is suppressed is thought to reflect the level of cognitive effort required by an upcoming task (reviewed in Van Diepen, Foxe, & Mazaheri, 2019). Greater alpha power suppression is associated with higher task demands (Fink et al., 2005), greater subjective task difficulty (Wostmann, Herrmann, Wilsch, & Obleser, 2015), increased memory load (Stipacek, Grabner, Neuper, Fink, & Neubauer, 2003), and greater discrimination difficulty (Roberts, Fedota, Buzzell, Parasuraman, & McDonald, 2014). Observing greater alpha suppression but no decrement in performance after inflammation induction in the current study suggests that inflammation heightens the cognitive effort needed to perform the task. This explanation seems in apparent contrast with the suggestion that motivational changes are a characteristic of inflammation (Felger & Treadway, 2016; Draper et al., 2017). However, careful inspection of the literature suggests that inflammation may result in a recalibration of reward-effort trade-offs, i.e., greater effort is invested when the behavior is regarded as especially rewarding (Lasselin et al., 2016; Vichaya, Hunt, & Dantzer, 2014) . Although the current task was not designed to assess reward-effort trade-offs, the observed pattern of results extends the notion of a reward-effort recalibration to attentional performance and suggests that increased effort is invested to maintain a high level of attentional performance. Arguing against this idea is the possibility that inflammation might degrade low-level visual sensory processing and thereby demand greater higher level processing that consequently leads to greater alpha power suppression (Roberts et al., 2014). However, such an explanation cannot account for effects based on difference scores between cue conditions. If inflammation degraded visual sensory processes, then greater alpha suppression would have been found for all cue conditions, resulting in no effect on attention difference scores.

In line with previous experimental studies using diverse tasks subsumed under attention, inflammation exhibited no overt behavioral attention effects (Brydon et al., 2008; Grigoleit et al., 2010; Krabbe et al., 2005; Reichenberg et al., 2001; van den Boogaard et al., 2010). However, absence of behavioral effects does not imply the absence of an underlying neurophysiological effect of inflammation. It has been shown that EEG can identify early signs of cognitive decline in pathological states, such as mild cognitive impairment (Mazaheri et al., 2018) and dementia in Parkinson’s disease (Bonanni et al., 2015; Klassen et al., 2011). EEG methods, as compared to most behavioral measures, are able to detect subtle aspects of cognitive function, making this method highly suitable for probing neural effects of mild inflammation. Indeed, the current results further underline the need for caution when drawing conclusions from non-significant behavioral results. Nevertheless, it remains unclear why behavior is unaffected by inflammation when underlying neurophysiological processes indicate significant alterations. One reason may be that overt behavioral effects of inflammation on attention require persistent (i.e., chronic) or severe inflammation before compensatory mechanisms that maintain performance fail to cope with weakened preparatory attention mechanisms. Typhoid vaccination, as used here, elicited only a 4-fold increase in IL-6 levels, whereas the endotoxin model, often used to experimentally study high inflammation, generally raises IL-6 levels 100-fold up to roughly 1000-fold (e.g., Draper et al., 2017; Eisenberger, Inagaki, Rameson, Mashal, & Irwin, 2009; Grigoleit et al., 2011; Lasselin et al., 2016; Muscatell et al., 2016). Yet such studies fail to show evidence supporting the possibility that inflammation affects overt attentional processes (reviewed in Bollen et al., 2017), suggesting that higher inflammation levels alone are not sufficient to induce overt attentional changes. The modest but reliable elevation of IL-6 observed in the current study is akin to the inflammation levels seen in subsets of depressed individuals, as well as in medical conditions such as diabetes and atherosclerosis (e.g., Dowlati et al., 2010; Maes et al., 1995; O’Brien, Scully, Fitzgerald, Scott, & Dinan, 2007). However, a difference between experimental models of inflammation and the aforementioned medical conditions is the duration of inflammation. Here it was up to 8 hours compared to the weeks, months or even years of elevated inflammation in these medical conditions. Considering that patients with chronic inflammation, such as those with cystic fibrosis and inflammatory bowel disease, show reduced attention performance as compared to healthy controls (Piasecki, Stanis, awska-Kubiak, Strzelecki, & Mojs, 2017) raises the possibility that overt behavioral effects of inflammation on attention only occur when inflammation is persistent. Perhaps, with chronic inflammation neural compensatory mechanisms eventually fail, allowing behavioral indices dependent on attentional preparation processes to become sensitive to inflammatory states.

The current findings suggest that EEG correlates of attention may be used to detect subtle neurophysiological changes accompanying inflammation. Future research may assess whether preparatory alpha suppression can be used as a pre-clinical predictive marker to identify those at risk to develop inflammation-associated changes. The current findings are also important in light of the high prevalence of mild cases of flu, colds or minor infections that cause mild, acute inflammation. Mild inflammatory states may enhance feelings of mental fatigue or cognitive stress due to the increase effort needed to perform otherwise effortless tasks
In sum, the present study showed, for the first time, a unique neurophysiological vulnerability of the alerting dimension of attention with transient mild inflammation. Future studies may explore the potential of neurophysiological outcomes as a marker for early detection of inflammation-associated cognitive changes.

## CONFLICT OF INTEREST

The authors declare no conflict of interests.

## ACKNOWLEDGEMENTS

The authors would like to thank Sasha Hulsken for her contribution in conceiving and executing the experiment, Farahdina Bachtiar, Anne Clemens, Jessica Maund, Alexandra Morrison, Greta Ontrup, and Marina Wissink for their valuable contribution in recruiting and testing participants and the nursing staff of the School of Nursing (University of Birmingham) for administering the injections.

## REFERENCES

Allison, D. J., & Ditor, D. S. (2014). The common inflammatory etiology of depression and cognitive impairment: a therapeutic target. Journal of Neuroinflammation, 11(1), 151. http://doi.org/10.1186/s12974-014-0151-1

Balter, L. J. T., Hulsken, S., Aldred, S., Drayson, M. T., Higgs, S., VeldhuijzenvanZanten, J. J. C. S., … Bosch, J. A. (2018). Low-grade inflammation decreases emotion recognition – Evidence from the vaccination model of inflammation. Brain, Behavior, and Immunity. http://doi.org/10.1016/j.bbi.2018.05.006

Bollen, J., Trick, L., Llewellyn, D., & Dickens, C. (2017). The effects of acute inflammation on cognitive functioning and emotional processing in humans: A systematic review of experimental studies. Journal of Psychosomatic Research. http://doi.org/10.1016/j.jpsychores.2017.01.002

Bonanni, L., Perfetti, B., Bifolchetti, S., Taylor, J. P., Franciotti, R., Parnetti, L., … Onofrj, M. (2015). Quantitative electroencephalogram utility in predicting conversion of mild cognitive impairment to dementia with Lewy bodies. Neurobiology of Aging. http://doi.org/10.1016/j.neurobiolaging.2014.07.009

Bourassa, K., & Sbarra, D. A. (2016). Body mass and cognitive decline are indirectly associated via inflammation among aging adults. Brain, Behavior, and Immunity. http://doi.org/10.1016/j.bbi.2016.09.023

Broadbent, D. (1966). Perception and communication. Education + Training. http://doi.org/10.1108/eb015727

Brydon, L., Harrison, N. A., Walker, C., Steptoe, A., & Critchley, H. D. (2008). Peripheral Inflammation is Associated with Altered Substantia Nigra Activity and Psychomotor Slowing in Humans. Biological Psychiatry, 63(11), 1022–1029. http://doi.org/10.1016/j.biopsych.2007.12.007

Capuron, L., Pagnoni, G., Demetrashvili, M., Woolwine, B. J., Nemeroff, C. B., Berns, G. S., & Miller, A. H. (2005). Anterior cingulate activation and error processing during interferon-alpha treatment. Biological Psychiatry, 58(3), 190–196. http://doi.org/10.1016/j.biopsych.2005.03.033

Cavanagh, J. F., & Frank, M. J. (2014). Frontal theta as a mechanism for cognitive control. Trends in Cognitive Sciences. http://doi.org/10.1016/j.tics.2014.04.012

Curran, S. L., Andrykowski, M. A., & Studts, J. L. (1995). Short Form of the Profile of Mood States (POMS-SF): Psychometric information. Psychological Assessment. http://doi.org/10.1037/1040-3590.7.1.80

Deckers, K., Schievink, S. H. J., Rodriquez, M. M. F., Van Oostenbrugge, R. J., VanBoxtel, M. P. J., Verhey, F. R. J., & Köhler, S. (2017). Coronary heart disease and risk for cognitive impairment or dementia: Systematic review and meta-analysis. PLoS ONE. http://doi.org/10.1371/journal.pone.0184244

Delorme, A., & Makeig, S. (2004). EEGLAB: An open source toolbox for analysis of single-trial EEG dynamics including independent component analysis. Journal of Neuroscience Methods, 134(1), 9–21. http://doi.org/10.1016/j.jneumeth.2003.10.009

Dhar, A. K., & Barton, D. A. (2016). Depression and the link with cardiovascular disease. Frontiers in Psychiatry. http://doi.org/10.3389/fpsyt.2016.00033

Dowlati, Y., Herrmann, N., Swardfager, W., Liu, H., Sham, L., Reim, E. K., & LanctÔt, K. L. (2010). A Meta-Analysis of Cytokines in Major Depression. Biological Psychiatry, 67(5), 446–457. http://doi.org/10.1016/j.biopsych.2009.09.033

Draper, A., Koch, R. M., van der Meer, J. W., Apps, M., Pickkers, P., Husain, M., & vander Schaaf, M. E. (2017). Effort but not Reward Sensitivity is Altered by Acute Sickness Induced by Experimental Endotoxemia in Humans. Neuropsychopharmacology, (August), 1–39. http://doi.org/10.1038/npp.2017.231

Eisenberger, N. I., Inagaki, T. K., Rameson, L. T., Mashal, N. M., & Irwin, M. R. (2009). An fMRI study of cytokine-induced depressed mood and social pain: The role of sex differences. NeuroImage, 47(3), 881–890. http://doi.org/10.1016/j.neuroimage.2009.04.040

Fan, J., Byrne, J., Worden, M. S., Guise, K. G., McCandliss, B. D., Fossella, J., & Posner, M. I. (2007). The Relation of Brain Oscillations to Attentional Networks. Journal of Neuroscience. http://doi.org/10.1523/JNEUROSCI.1833-07.2007

Fan, J., McCandliss, B. D., Sommer, T., Raz, A., & Posner, M. I. (2002). Testing the efficiency and independence of attentional networks. Journal of Cognitive Neuroscience, 14(3), 340–347. http://doi.org/10.1162/089892902317361886

Felger, J. C., & Treadway, M. T. (2016). Inflammation Effects on Motivation and Motor Activity: Role of Dopamine. Neuropsychopharmacology: Official Publication of the American College of Neuropsychopharmacology, 42 (August), 1–88. http://doi.org/10.1038/npp.2016.143

Fink, A., Grabner, R. H., Neuper, C., & Neubauer, A. C. (2005). EEG alpha band dissociation with increasing task demands. Cognitive Brain Research. http://doi.org/10.1016/j.cogbrainres.2005.02.002

Freed, R. D., Mehra, L. M., Laor, D., Patel, M., Alonso, C. M., Kim-Schulze, S., & Gabbay, V. (2018). Anhedonia as a clinical correlate of inflammation in adolescents across psychiatric conditions. World Journal of Biological Psychiatry. http://doi.org/10.1080/15622975.2018.1482000

Grigoleit, J. S., Kullmann, J. S., Wolf, O. T., Hammes, F., Wegner, A., Jablonowski, S., … Schedlowski, M. (2011). Dose-dependent effects of endotoxin on neurobehavioral functions in humans. PLoS ONE, 6(12). http://doi.org/10.1371/journal.pone.0028330

Grigoleit, J. S., Oberbeck, J. R., Lichte, P., Kobbe, P., Wolf, O. T., Montag, T., … Schedlowski, M. (2010). Lipopolysaccharide-induced experimental immune activation does not impair memory functions in humans. Neurobiology of Learning and Memory. http://doi.org/10.1016/j.nlm.2010.09.011

Haegens, S., Handel, B. F., & Jensen, O. (2011). Top-Down Controlled Alpha Band Activity in Somatosensory Areas Determines Behavioral Performance in a Discrimination Task. Journal of Neuroscience. http://doi.org/10.1523/jneurosci.5199-10.2011

Harrison, N. A., Voon, V., Cercignani, M., Cooper, E. A., Pessiglione, M., & Critchley, H. D. (2015). A Neurocomputational Account of How Inflammation Enhances Sensitivity to Punishments Versus Rewards. Biological Psychiatry. http://doi.org/10.1016/j.biopsych.2015.07.018

Keil, A., Mussweiler, T., & Epstude, K. (2006). Alpha-band activity reflects reduction of mental effort in a comparison task: A source space analysis. Brain Research. http://doi.org/10.1016/j.brainres.2006.08.118

Klassen, B. T., Hentz, J. G., Shill, H. A., Driver-Dunckley, E., Evidente, V. G. H., Sabbagh, M. N., … Caviness, J. N. (2011). Quantitative EEG as a predictive biomarker for Parkinson disease dementia. Neurology. http://doi.org/10.1212/WNL.0b013e318224af8d

Klimesch, W., Sauseng, P., & Hanslmayr, S. (2007). EEG alpha oscillations: The inhibition-timing hypothesis. Brain Research Reviews. http://doi.org/10.1016/j.brainresrev.2006.06.003

Krabbe, K. S., Reichenberg, A., Yirmiya, R., Smed, A., Pedersen, B. K., & Bruunsgaard, H. (2005). Low-dose endotoxemia and human neuropsychological functions. Brain, Behavior, and Immunity, 19(5), 453–460. http://doi.org/10.1016/j.bbi.2005.04.010

Kullmann, J. S., Grigoleit, J.-S., Wolf, O. T., Engler, H., Oberbeck, R., Elsenbruch, S., … Gizewski, E. R. (2013). Experimental human endotoxemia enhances brain activity during social cognition. Social Cognitive and Affective Neuroscience. http://doi.org/10.1093/scan/nst049

Kurella Tamura, M., Tam, K., Vittinghoff, E., Raj, D., Sozio, S. M., Rosas, S. E., … Townsend, R. R. (2017). Inflammatory Markers and Risk for Cognitive Decline in Chronic Kidney Disease: The CRIC Study. Kidney International Reports. http://doi.org/10.1016/j.ekir.2016.10.007

Lasselin, J., Treadway, M. T., Lacourt, T. E., Soop, A., Olsson, M. J., Karshikoff, B., … Lekander, M. (2016). Lipopolysaccharide Alters Motivated Behavior in a Monetary Reward Task: a Randomized Trial. Neuropsychopharmacology, 1–10. http://doi.org/10.1038/npp.2016.191

Lin, T., Liu, G. A., Perez, E., Rainer, R. D., Febo, M., Cruz-Almeida, Y., & Ebner, N. C. (2018). Systemic Inflammation Mediates Age-Related Cognitive Deficits. Frontiers in Aging Neuroscience,10(August), 1–9. http://doi.org/10.3389/fnagi.2018.00236

Luppino, F. S., De Wit, L. M., Bouvy, P. F., Stijnen, T., Cuijpers, P., Penninx, B. W. J. H., & Zitman, F. G. (2010). Overweight, obesity, and depression: A systematic review and meta-analysis of longitudinal studies. Archives of General Psychiatry. http://doi.org/10.1001/archgenpsychiatry.2010.2

Maes, M., Meltzer, H. Y., Bosmans, E., Bergmans, R., Vandoolaeghe, E., Ranjan, R., & Desnyder, R. (1995). Increased plasma concentrations of interleukin-6, soluble interleukin-6, soluble interleukin-2 and transferrin receptor in major depression. Journal of Affective Disorders, 34(4), 301–309. http://doi.org/10.1016/0165-032700028-L(95)

Maris, E., & Oostenveld, R. (2007). Nonparametric statistical testing of EEG-and MEG-data.Journal of Neuroscience Methods. http://doi.org/10.1016/j.jneumeth.2007.03.024

Marsland, A. L., Petersen, K. L., Sathanoori, R., Muldoon, M. F., Neumann, S. A., Ryan, C., … Manuck, S. B. (2006). Interleukin-6 covaries inversely with cognitive performance among middle-aged community volunteers. Psychosomatic Medicine, 68(6), 895–903. http://doi.org/10.1097/01.psy.0000238451.22174.92

Mazaheri, A., Segaert, K., Olichney, J., Yang, J. C., Niu, Y. Q., Shapiro, K., & Bowman, H. (2018). EEG oscillations during word processing predict MCI conversion to Alzheimer’s disease. NeuroImage: Clinical. http://doi.org/10.1016/j.nicl.2017.10.009

Muscatell, K. A., Moieni, M., Inagaki, T. K., Dutcher, J. M., Jevtic, I., Breen, E. C., … Eisenberger, N. I. (2016). Exposure to an inflammatory challenge enhances neural sensitivity to negative and positive social feedback. Brain, Behavior, and Immunity, 57, 21–29. http://doi.org/10.1016/j.bbi.2016.03.022

O’Brien, S. M., Scully, P., Fitzgerald, P., Scott, L. V., & Dinan, T. G. (2007). Plasma cytokine profiles in depressed patients who fail to respond to selective serotonin reuptake inhibitor therapy. Journal of Psychiatric Research, 41(3–4), 326–331. http://doi.org/10.1016/j.jpsychires.2006.05.013

Oostenveld, R., Fries, P., Maris, E., & Schoffelen, J. M. (2011). FieldTrip: Open source software for advanced analysis of MEG, EEG, and invasive electrophysiological data. Computational Intelligence and Neuroscience, 2011. http://doi.org/10.1155/2011/156869

Paine, N. J., Ring, C., Bosch, J. A., Drayson, M. T., & Veldhuijzen vanZanten, J. J. C. S. (2013). The time course of the inflammatory response to the Salmonella typhi vaccination. Brain, Behavior, and Immunity, 30, 73–9. http://doi.org/10.1016/j.bbi.2013.01.004

Piasecki, B., StanisŁawska-Kubiak, M., Strzelecki, W., & Mojs, E. (2017). Attention and memory impairments in pediatric patients with cystic fibrosis and inflammatory bowel disease in comparison to healthy controls. Journal of Investigative Medicine: The Official Publication of the American Federation for Clinical Research, 65(7), 1062–1067. http://doi.org/10.1136/jim-2017-000486

Posner, M. I., & Rothbart, M. K. (2007). Research on Attention Networks as a Model for the Integration of Psychological Science. Annual Review of Psychology, 58(1), 1–23. http://doi.org/10.1146/annurev.psych.58.110405.085516

Reichenberg, A., Yirmiya, R., Schuld, A., Kraus, T., Haack, M., Morag, A., & Pollmächer, T. (2001). Cytokine-associated emotional and cognitive disturbances in humans. Archives of general psychiatry (Vol. 58).

Roberts, D. M., Fedota, J. R., Buzzell, G. A., Parasuraman, R., & McDonald, C. G. (2014). Prestimulus Oscillations in the Alpha Band of the EEG Are Modulated by the Difficulty of Feature Discrimination and Predict Activation of a Sensory Discrimination Process. Journal of Cognitive Neuroscience. http://doi.org/10.1162/jocn_a_00569

Sawaki, R., Luck, S. J., & Raymond, J. E. (2015). How attention changes in response to incentives. Journal of Cognitive Neuroscience. http://doi.org/10.1162/jocn_a_00847

Singh-Manoux, A., Dugravot, A., Brunner, E., Kumari, M., Shipley, M., Elbaz, A., & Kivimaki, M. (2014). Interleukin-6 and C-reactive protein as predictors of cognitive decline in late midlife. Neurology. http://doi.org/10.1212/WNL.0000000000000665

Smith, A. P. (2012). Effects of the common cold on mood, psychomotor performance, the encoding of new information, speed of working memory and semantic processing. Brain, Behavior, and Immunity, 26(7), 1072–1076. http://doi.org/10.1016/j.bbi.2012.06.012

Stipacek, A., Grabner, R. H., Neuper, C., Fink, A., & Neubauer, A. C. (2003). Sensitivity of human EEG alpha band desynchronization to different working memory components and increasing levels of memory load. Neuroscience Letters. http://doi.org/10.1016/j.neulet.2003.09.044

van den Boogaard, M., Ramakers, B. P., van Alfen, N., van der Werf, S. P., Fick, W. F., Hoedemaekers, C. W., … Pickkers, P. (2010). Endotoxemia-induced inflammation and the effect on the human brain. Critical Care (London, England), 14(3), R81. http://doi.org/10.1186/cc9001

Van Diepen, R., Foxe, J. J., & Mazaheri, A. (2019). The functional role of alpha-band activity in attentional processing: The current zeitgeist and future outlook. Current Opinion in Psychology. http://doi.org/10.1016/J.COPSYC.2019.03.015

Vichaya, E. G., Hunt, S. C., & Dantzer, R. (2014). Lipopolysaccharide Reduces Incentive Motivation While Boosting Preference for High Reward in Mice. Neuropsychopharmacology, 39(10), 2884–2890. http://doi.org/10.1038/npp.2014.141

Vollmer-Conna, U. acute;, Fazou, C., Cameron, B., Li, H., Brennan, C., Luck, L., … Lloyd, A. (2004). Production of pro-inflammatory cytokines correlates with the symptoms of acute sickness behaviour in humans. Psychological Medicine. http://doi.org/10.1017/S0033291704001953

Wagenmakers, E.-J., Love, J., Marsman, M., Jamil, T., Ly, A., Verhagen, J., … Morey, R. D. (2017). Bayesian inference for psychology. Part II: Example applications with JASP. Psychonomic Bulletin & Review. http://doi.org/10.3758/s13423-017-1323-7

Wostmann, M., Herrmann, B., Wilsch, A., & Obleser, J. (2015). Neural Alpha Dynamics in Younger and Older Listeners Reflect Acoustic Challenges and Predictive Benefits. Journal of Neuroscience. http://doi.org/10.1523/JNEUROSCI.3250-14.2015

